# KofamKOALA: KEGG ortholog assignment based on profile HMM and adaptive score threshold

**DOI:** 10.1101/602110

**Authors:** Takuya Aramaki, Romain Blanc-Mathieu, Hisashi Endo, Koichi Ohkubo, Minoru Kanehisa, Susumu Goto, Hiroyuki Ogata

**Affiliations:** Institute for Chemical Research, Kyoto University, Gokasho, Uji, Kyoto 611-0011, Japan; Hewlett-Packard Japan, Ltd. 2-2-1, Ojima, Koto-ku, Tokyo 136-8711, Japan; Database Center for Life Science, Research Organization of Information and Systems, Kashiwa, Chiba 277-0871, Japan

## Abstract

**Summary:** KofamKOALA is a web server to assign KEGG Orthologs (KOs) to protein sequences by homology search against a database of profile hidden Markov models (KOfam) with pre-computed adaptive score thresholds. KofamKOALA is faster than existing KO assignment tools with its accuracy being comparable to the best performing tools. Function annotation by KofamKOALA helps linking genes to KEGG resources such as the KEGG pathway maps and facilitates molecular network reconstruction.

**Availability:** KofamKOALA, KofamScan, and KOfam are freely available from https://www.genome.jp/tools/kofamkoala/

## 1 Introduction

Automatic gene function annotation is an important first step to interpret genomic data. KEGG (Kyoto Encyclopedia of Genes and Genomes) is a widely used reference knowledge base, which helps investigate genomic functions by linking genes to biological knowledge such as metabolic pathways and molecular networks (1). In KEGG, the KEGG Orthology (KO) database − a manually curated large collection of protein families − serves as a baseline reference to link genes with other KEGG resources such as metabolic maps. Currently, KO identifiers (i.e., K numbers) are assigned to 12,934,525 (48%) protein sequences in the KEGG GENES database (27,173,868 proteins).

Three existing tools, BlastKOALA, GhostKOALA (2) and KAAS (3), are currently available to assign KOs to protein sequences. These tools use homology search software such as BLAST (4) and GHOSTX (5) to search amino acid sequences against GENES. To reduce large computational times required for multiple pairwise sequence comparisons, these tools use a subset of representative sequences in GENES to build their target database. In this study, we propose to employ profile hidden Markov model (HMM) to compress the database and to define adaptive thresholds for similarity scores to reliably assign K numbers to protein sequences.

## 2 Implementation

For each group of orthologous protein sequences in GENES annotated with a given KO (K number), we generate a profile HMM in the following way. First, sequence redundancy in the sequence set is reduced by CD-HIT (6) with 100% sequence identity clustering cutoff. Next, MAFFT (7) and HMMER/hmmbuild (8) are used to align sequences and to generate a profile HMM.

An adaptive score threshold is computed for each HMM in the following way. Sequence similarity score (bit score) between a protein sequence and an HMM is computed using HMMER/hmmsearch. The non-redundant sequences belonging to the corresponding KO family are divided into three groups. One of the groups is used as the positive dataset, while the sequences in the remaining two groups are used to generate a profile HMM. Sequences belonging to other KO families serve as the negative dataset for the KO in consideration. Based on the set of bit scores between the profile HMM and the sequences in the positive/negative datasets, we determine a threshold score, *T*, to maximize the *F*-measure [where *F* = 2/(*Recall*^−1^ + *Precision*^−1^)]. This procedure is repeated three times by replacing the positive dataset among the three groups and the average of *T* 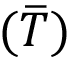 is defined as the adaptive threshold score for the assignment of the K-number to a sequence.

The database of HMMs with score thresholds was named KOfam. We developed KofamScan and KofamKOALA to annotate genes using KOfam and to link them with other KEGG resources for versatile functional investigation. The former is a command line script, while the latter is a web implementation of the script and the database.

## 3 Assessment

To compare the performance of KofamScan with BlastKOALA, GhostKOALA and KAAS, we used 40 genomes (20 eukaryotes and 20 prokaryotes; Supplementary Table S1) randomly selected from the GENES database as test queries. This test set contains 383,202 sequences (143,662 sequences with K-number assignment) corresponding to 16,166 distinct K-numbers. From the GENES database, we removed all the genomes belonging to the genera selected as test queries. Then, using the remining GENES sequences annotated with K-numbers, we generated a test KOfam database for this assessment. As for BlastKOALA, GhostKOALA and KAAS, we used the default target databases used in their respective web servers after removing genomes from the genera that we selected for test queries.

The KOfam database created for this test assessment contained 20,394 profile HMMs, of which 9,414 KOs were represented by prokaryotic sequences. For the 40 genomes constituting our test set, prediction accuracy (*F*-measure) was comparable among KofamScan (0.865), BlastKOALA (0.888), and GhostKOALA (0.861), while KAAS showed a lower *F*-measure (0.809) (Fig. 1). To perform another test using only 20 prokaryotic genomes as test queries, we reduced the target databases either by excluding profile HMMs composed exclusively of eukaryotic sequences (for KofamScan) or by using the target database for prokaryotes (for BlastKOALA, GhostKOALA and KAAS). Again, the prediction accuracy of KofamScan (*F*=0.875) was comparable to BlastKOALA (0.846), GhostKOALA (0.886), and KAAS showed a lower accuracy (0.786) (Fig. 1).

**Fig. 1.**
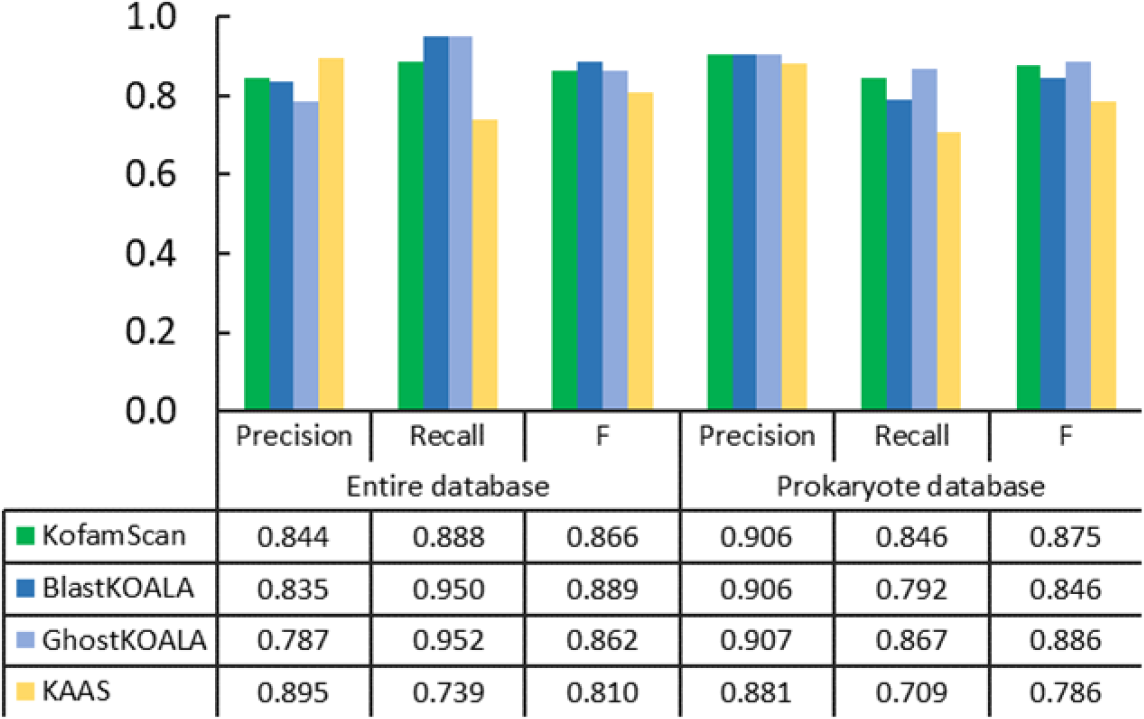
Comparison of the performance of KofamScan with other tools.

Regarding computational speed, KofamScan was 69 times faster than BlastKOALA, whereas GhostKOALA and KAAS were respectively 65 and 33 times faster than BlastKOALA for the test with 40 genomes (Supplementary Table S2). For the test with 20 prokaryote genomes, KofamScan was 83 times faster than BlastKOALA, whereas GhostKOALA and KAAS were 47 and 42 times faster than BlastKOALA, respectively. Therefore, the effect of the reduction of the target database size is more pronounced for KofamScan compared to the three other tools while conserving amongst the highest levels of prediction accuracy.

## 4 Summary

We developed a database of profile HMMs based on the KO and GENES databases. The adaptive score thresholds are precalculated for individual KO families, and can be used to assign KO to sequences using KofamScan and KofamKOALA. Sequence matches with a score exceeding the score threshold are considered more reliable than other matches and highlighted with ‘*’ marks in the output of these tools. The web implemented KofamKOALA tool has additional functions to automatically send the search results to KEGG Mapper for reconstruction of pathways (PATHWAY), pathway modules (MODULE) and hierarchical function classifications (BRITE). KofamScan and KOfam can be downloaded freely from the GenomeNet FTP server (ftp://ftp.genome.jp/). Users may be able to customize the KOfam database by selecting KOs of interest, so that they can focus on specific protein functions for their studies.

## Acknowledgements

Computation time was provided by the SuperComputer System, Institute for Chemical Research, Kyoto University.

## Funding

This work has been supported by JSPS/MEXT/KAKENHI (Nos. 26430184, 18H02279, 16H06429, 16K21723, 16H06437) and the Collaborative Research Program of the Institute for Chemical Research, Kyoto University (No. 2018-30).

## Conflict of Interest

none declared.

