## Supplementary data for "KofamKOALA: KEGG ortholog assignment based on profile HMM and adaptive score threshold"

**Details of implementation**

For each set of protein sequences with a same KO (K number), we generate a profile hidden Markov model (HMM) in the following way. First, sequence redundancy in the set of sequences is reduced by CD-HIT version 4.7 (1) with 100% sequence identity cutoff to produce a set of non-redundant sequences. Next, MAFFT version 7.310 (2) is used (options, --amino --anysymbol) to align sequences. Finally, HMMER/hmmbuild ver. 3.1b2 (3) is used to build a profile HMM from the alignment.

An adaptive score threshold is computed for each HMM in the following way. Sequence similarity score (bit score) between a protein sequence and an HMM is computed using HMMER/hmmsearch. The non-redundant sequences belonging to the corresponding KO family are divided into three groups of equal sizes. A group is used as the positive dataset, while the sequences in the remaining two groups are used to generate a profile HMM by the above described procedure. Sequences belonging to other KO families serve as the negative dataset for the KO under consideration. Based on the set of bit scores between the HMM and the sequences in the positive/negative datasets, we determine a threshold score, $T$, that maximizes the $F$-measure [defined as $F = 2/(Recall^{-1} + Precision^{-1})$]. This procedure is repeated three times by replacing the positive dataset among the three groups. Then, we obtain the average of $T$ (denoted by $\bar{T}$) and the average of $F$ (denoted by $\bar{F}$). $\bar{T}$ is defined as the adaptive threshold score for the corresponding profile HMM to assign its K-number to query sequences.

We obtain two types of bit score by hmmsearch (i.e., "the full sequence bit score" and "the best domain bit score"). In addition, alignments are used either as they are produced by the above described procedure ("raw") or with an additional processing by trimAl (4) (with options, -resoverlap 75 -seqoverlap 0.75) to remove distantly related sequences in the alignments ("trimmed"). Through the combination of the two types of bit score ("full score" or "domain score") and two types of alignment ("raw" or "trimmed"), we obtain four values for the adaptive threshold $\bar{T}$ and for values for the associated $\bar{F}$ for each KO (**Figure S1**). Based on $\bar{F}$ for individual combinations, we select the best performing combination of score and alignment types for each KO. The selected profile HMMs are stored in the resulting profile HMM database, named KOfam, with associated $\bar{T}$ and $\bar{F}$.

KofamScan is a Ruby script that utilizes hmmsearch, KOfam and the adaptive thresholds for function annotation. In cases where $\bar{T}$ could not be generated due to small number of sequences (less than three) in individual KO, we set the common threshold for these KOs by taking the average of $\bar{T}$ across different KOs.

KofamKOALA is a web implementation of KofamScan to search against KOfam, with additional functions to automatically send the KofamScan results to KEGG Mapper for reconstruction of pathways (PATHWAY), pathway modules (MODULE) and hierarchical function classifications (BRITE).

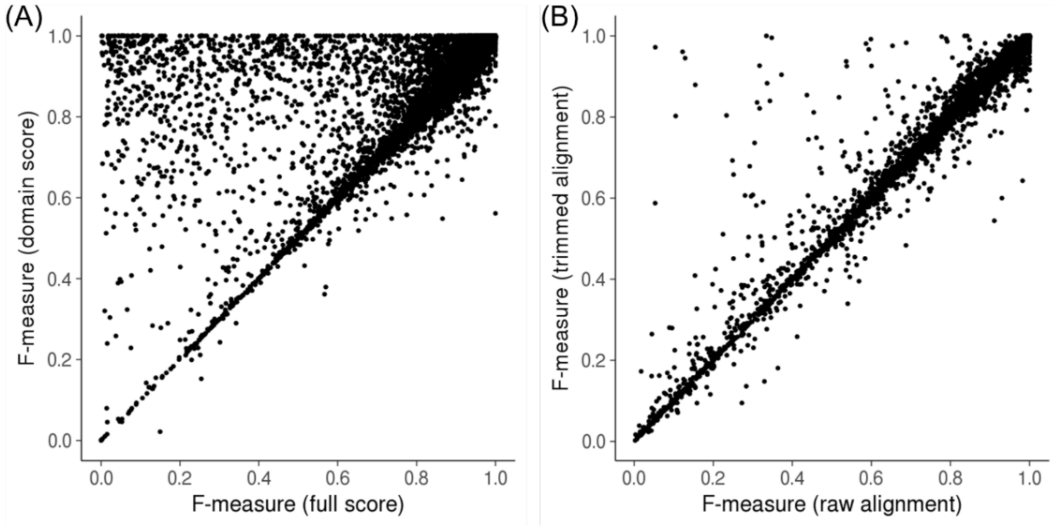


**Figure S1.** Effect of types of score and alignment processing. (A) Comparison of F-measures between “the full sequence bit score (raw alignment)” and “the best domain bit score (raw alignment)”. (B) Comparison of *F*-measures between raw and trimmed alignments (full sequence bit score).

**Performance assessment**

To compare KofamScan with BlastKOALA, GhostKOALA (5) and KAAS (6), we randomly selected 40 genomes (20 eukaryotes and 20 prokaryotes; **Table S1**) from the KEGG GENES database (release 88.2) and used these genomes as test queries. From the GENES database, we removed all the genomes belonging to the genera that were selected for the test queries. Then, using the remaining GENES sequences and the KO database, we generated a KOfam database for this test. In this test, we defined *Recall* and *Precision* as follows:

$$Recall = \frac{match}{match + unmatch + missing}$$

$$Precision = \frac{match}{match + unmatch + excess}$$

In these equations, *match* is the number of cases where predicted KO is identical to the KO defined in GENES; *unmatch* is the number of cases where predicted KO is different from the KO defined in GENES; *missing* is the number of cases where KO is defined in GENES but no prediction was made; *excess* is the number of cases where KO is not defined in GENES, but prediction assigned one or more KOs. *F*-measure was computed by $F = 2/(Recall^{-1} + Precision^{-1})$. The results of the comparative assessment are given in **Figure 1** in the main manuscript as well as in **Table S2**.

**Table S1.** Genomes used for the comparative assessment of KofamScan with three other tools.

| | |
|---|---|
| Eukaryotes | *Ailuropoda melanoleuca* (aml), *Apis mellifera* (ame), *Aspergillus fumigatus* (afm), *Aspergillus nidulans* (ani), *Aspergillus oryzae* (aor), *Bos taurus* (bta), *Brugia malayi* (bmy), *Candida glabrata* (cgr), *Drosophila melanogaster* (dme), *Leishmania infantum* (lif), *Malassezia globosa* (mgl), *Mus musculus* (mmu), *Nematostella vectensis* (nve), *Ornithorhynchus anatinus* (oaa), *Plasmodium knowlesi* (pkn), *Pyricularia oryzae* 70-15 (mgr), *Strongylocentrotus purpuratus* (spu), *Thalassiosira pseudonana* (tps), *Toxoplasma gondii* (tgo), *Trichomonas vaginalis* (tva) |
| Prokaryotes | *Alicycliphilus denitrificans* BC (and), *Bacillus licheniformis* ATCC 14580 (bli), *Bordetella avium* (bav), *Borrelia afzelii* (baf), *Borrelia duttonii* (bdu), *Burkholderia* sp. CCGE1002 (bge), *Comamonas thiooxydans* (ctt), *Deinococcus radiodurans* (dra), *Geobacter daltonii* FRC-32 (geo), *Halorhodospira halophila* (hha), *Leuconostoc mesenteroides* subsp. *mesenteroides* ATCC 8293 (lme), *Methanoregula boonei* (mbn), *Pseudonocardia dioxanivorans* (pdx), *Pyrococcus horikoshii* (pho), *Rhodobacter sphaeroides* KD131 (rsk), *Saccharophagus degradans* (sde), *Synechococcus* sp. CC9902 (sye), *Thermotoga* sp. RQ2 (trq), *Teredinibacter turnerae* (ttu), *Tropheryma whipplei* Twist (twh) |

Species name is followed by the KEGG organism code in parentheses.

**Table S2.** CPU times required for the tests.

| | Entire target database and 40 query genomes | | Reduced target database and 20 query prokaryotic genomes | |
|---|---|---|---|---|
| | CPU time | Speed ratio | CPU time | Speed ratio |
| KofamScan | 2h26m18s | 68.99 | 0h11m59s | 82.80 |
| GhostKOALA | 2h34m50s | 65.19 | 0h21m02s | 47.15 |
| KAAS | 5h03m32s | 33.25 | 0h23m11s | 42.80 |
| BlastKOALA | 168h12m41s | 1.00 | 16h31m53s | 1.00 |

*Res* 35(Web Server issue):W182-185.
